# Symbolic and Non-Symbolic numbers differently affect center identification in a number-line bisection task

**DOI:** 10.1101/2024.12.01.626261

**Authors:** Annamaria Porru, Lucia Ronconi, Daniela Lucangeli, Lucia Regolin, Silvia Benavides-Varela, Rosa Rugani

## Abstract

Numerical and spatial representations are intertwined as in the Mental Number Line, where smaller numbers are on the left and larger numbers on the right. This relationship has been repeatedly demonstrated with various experimental approaches, such as the line bisection task.

Spatial accuracy appears to be systematically distorted leftward for smaller digits by elaboration of spatial codes during number processing. Other studies have investigated perceptual and visuo-spatial attention bias using the digit line bisection task, suggesting that these effects may be related to a cognitive illusion in which the reference numbers project their values onto the straight line, creating an illusory lateral disparity. On the other hand, both dot arrays (non-symbolic stimuli) and arabic numbers (symbolic stimuli) demonstrate a privileged relation between spatial and numerical elaboration. The bias toward the larger numerosity flanker was attributed to a length illusion. There is, however, no consensus regarding whether physical features and symbolic and non-symbolic numerical representations exert the same influence over spatial ones.

In the present study, we carried out a series of 4 Experiments to provide further evidence for a better understanding of the nature of this differential influence. All experiments presented the numbers in both symbolic and non-symbolic formats. In Experiment 1, the numbers “2-8” were presented in a variety of left-right orientations. In Experiment 2, the flankers were identical, “2-2” or “8-8”, and symmetrically displaced with respect to the line. In Experiment 3, we employed asymmetrically distributed eight dots, or font sizes in “8-8” numerals, to create a perceptual imbalance. In Experiment 4, we replicated the manipulation used in Experiment 3, but with two dots and “2-2” numerals.

The Non-Symbolic format induced stronger leftward biases, particularly when the larger numerosity (Experiment 1) or the denser stimuli near the line (Experiments 3 and 4) were on the left, while no bias emerged when flankers were numerically equivalent and symmetrical (Experiment 2). The left bias may result from a tendency to estimate the influence of stimulus perception associated with participant’ scanning direction, similar to the direction of pseudoneglect. Conversely, the Symbolic format induced mostly right bias, possibly due to left-lateralized processing and a tendency to use a common strategy involving scanning from left to right.

Altogether our data support the view that abstract numbers and non-symbolic magnitude affect perceptual and attentional biases, yet in distinctive ways.

## Introduction

Humans organize numerical information in space according to a specific left-to-right orientation: the so-called Mental Number Line (MNL; 1). The evidence for the orientation of the mental number line representation comes from the SNARC effect (Spatial-Numerical Association of Response Codes), which has been interpreted on the basis of a spatial-congruency between the response side (the left and right sides of egocentric space), and the relative position of the represented numerical magnitude on an oriented mental number line (the left-space/small numbers and the right space/large numbers) (2,3). More generally, a tight space and number association (SNA) can be observed in varied tasks, age groups, and even species (4).

For example, to investigate the influence of number magnitude on spatial attention, Martin Fischer (5) designed a paper-and-pencil task requiring neurologically healthy participants to mark the midpoint of strings composed of Arabic numerals. Strings formed by 1’s or 2’s induced a left bias, while strings of 8’s or 9’s induced a right bias. When strings were replaced with lines flanked by numerals, the midpoint was misjudged toward the flanker representing the larger magnitude (5). Such biases have been replicated when one of the flankers were within (i.e. 2-9) or outside (i.e. 5-8) the subitizing range (6). Moreover the bias is more pronounced with number words (e.g. ‘DEUX’, ‘NEUF’) than numerals (e.g. 2, 9) (7), and with larger numerical intervals (e.g. 1-8, 2-9) than smaller ones (e.g. 1-2, 8-9) (8). When assessed with this task, neglect patients showed a right bias when lines were flanked by large numerals (9-9) and a left bias when the flankers were small (1-1) (9). Altogether these studies provide evidence that numerical information systematically bias spatial performance in the line bisection task, supporting the hypothesis of an automatic association between numbers and space (5).

Perceptual and visuo-spatial attentional biases have been also documented when participants are presented with Arabic digits in the line bisection task. In the first experiment of a series, De Hevia and colleagues (10) used a bisection task with a string of numbers, and contrary to Fischer’s (5) findings, the authors did not find a number-dependent effect, but rather an effect that depended on the position of the string on the sheet. The second study used a line bisection task with numbers as flankers. Again, the numbers did not modulate the effect, but a tendency to bisect the line to the left which was attributed to a pseudoneglect phenomenon. In the third experiment, a bisection task was presented without the line, with the numbers physically positioned near or far. Once more, the number did not appear to modulate the effect, but it appears that the physical proximity of the numbers made it easier to read the numerical magnitudes. In the final experiment, participants were asked to bisect the lines or spaces between the two numbers in two different conditions, one in which they had to name the small number and one in which they had to name the large number. In this case, a bias toward the larger number was only found in the condition in which participants were asked to name the larger number, demonstrating a visual spatial orientation effect of attention. Therefore, the authors conclude that the effect found may be related to a cognitive illusion in which the numerical cues create an illusory lateral disparity (10).

A subsequent implementation of a non-symbolic version of the task, with numerical information presented in the form of dot arrays, allowed testing the same task in adults, primary school, and preschool children (11). Results showed that, whereas a spatial bias with numerals manifests only in adults, the non-symbolic format (arrays of 2 or 9 dots) systematically distorted the localization of the midpoint of a horizontal line at all three ages. Because adults showed a consistent bias toward the more numerous array in both the symbolic and the non-symbolic format (11), this was taken as evidence that the underlying numerical representation is common to both formats. This interpretation is also in agreement with other behavioral and brain imaging studies carried out with the comparison task (e.g., 12–14) and consistent with the hypothesis of a generalized magnitude system that represents different magnitudes (e.g., size, quantity, number) at different levels of abstraction (15) in an integrated manner (16). Moreover, the numerical bias in line bisection tasks, is thought to be independent of the format, and thus reflects an abstract mapping between numerical magnitude and horizontal space (11; for consistent results see 17–19).

Nevertheless, contrasting evidence has been also documented. Increasing number of findings support a distinction between symbolic and non-symbolic numbers’ representations (20–27), questioning the conclusion that the spatial biases observed with either dot-arrays or Arabic numbers share a common mechanism. In the line bisection tasks, specifically, the visuo-perceptual features of the dot-arrays appear to play a crucial role in determining the direction of the biases. For example, Gebuis and Gevers (28) showed that participants identified the midpoint closer when larger numerosity had the smallest total area. Additionally, when flankers were numerically identical but varied in area or surface, participants consistently bisected toward the larger area (6), highlighting the modulating role of perceptual cues in numerosity processing. Similarly, in another study, when symbolic and non-symbolic numbers were simultaneously displayed (with digits presented instead of dots, in a dice-like pattern), the automatic association between numbers and space emerged exclusively for the symbolic format, thereby reinforcing the concept of distinct representations for the two formats (20).

At present, this mixed evidence leaves open the question of whether symbolic and non-symbolic formats induce comparable effects on the processing of spatial extension in line bisection tasks. Here, we address this question by specifically exploring whether unbalancing different perceptual properties of the flankers in the two formats influences the core aspects of the automatic association between numbers and space to the same extent.

In Experiment 1, which was inspired by the study of De Hevia & Spelke (11), lines were flanked by different numbers (2-8 or 8-2) that were presented as either Arabic digits (Symbolic format) or dots (Non-Symbolic format). Under the hypothesis of shared numerical representations, participants were expected to show similar biases regardless of the format. In the following experiments, instead, lines were flanked by numerically identical flankers. In Experiment 2 the flankers, independently of the format, were physically exactly the same and symmetrically positioned relative to the line. We expected either an accurate middle identification due to the symmetry of the display and equal influence of the symbolic and non-symbolic flankers on the line, or alternatively, a leftward bias, consistent with the findings reported by De Hevia et al. (10). In Experiments 3 and 4, the perceptual saliency of the left and right flankers differed. In the Symbolic format, one digit had a larger font size. In the Non-Symbolic format, there was an overt unbalanced distribution of dots on the left and right half of the display, such that many dots were placed near the line and a few far from it, or *viceversa*. The latter experiments aimed at disentangling the influence of numerosity from that of perceptual features. Indeed, we have considered the possibility that salience and spatial cues could influence bisection when we manipulated the flankers. If the abstract numerical information extracted from the array is crucial, and the format and perceptual magnitude are irrelevant, no bias is expected. Alternatively, if the more salient stimulus is prevalent, we expect bias toward it.

Additionally, we addressed the question of whether the orientation of the number from left-to-right (Small-left) or right-to-left (Large-left) could produce comparable effects in the processing of spatial extent in line bisection tasks. In Experiment 1, based on De Hevia et al, (11), we expected participants to show a bias towards the larger number, regardless of orientation. In Experiment 2, due to the symmetry of the display and the perceptual balance between the two orientations, we expected either precise middle identification or a bias towards the left, as observed by De Hevia et al. (10). In Experiments 3 and 4, where the perceptual salience of the left and right flankers varied, we expected a bisection towards the flanker denser in the line proximity, consistent with the hypothesis that the more salient flanker would influence the bisection point more.

## Method

### Participants

An a priori power analysis was conducted using G*Power 3 icon (Version 3.1.9.6) to determine the appropriate sample size for our study. Based on the study by De Hevia and Spelke (11), we set the following parameters: power = .80, α = .05, Effect size Cohen’s f = .25. The power analysis was originally designed for repeated-measures within-subjects ANOVA. However, the actual analysis conducted in this study employed Linear Mixed Effects Models (LMMs), which better accounts for individual differences and the complex structure of the data.

The parameters selected for the ANOVA offer a general estimation. Given that LMMs frequently provide enhanced flexibility and power in detecting effects due to their capacity to model random effects, we consider the power analysis conducted here to be a conservative estimation.

According to the results of the power analysis, the total sample size needed is 34 participants for each experiment.

This study comprised 4 different groups. Thus, in total 136 participants were enrolled, 34 in each group. Ten participants were excluded during the analyses because their mean bias was more than 2 SD above the group mean. Three participants were replaced in each of Experiments 1, 2, and 3, and one in Experiment 4. For this reason, we recruited an additional 10 more participants to reach the planned sample size. All participants were Italian students at the University of Padua, Italy. The mean age was 19.89 years (SD = 1.35). All participants were right-handed and had normal or corrected-to-normal sight. All participant provided signed written informed consent prior to the study and were naïve about the hypotheses of the experiment. Participants were recruited between May 28 and July 28, 2023. The numerical bias phenomenon was explained at the end of the experiment.

This study was approved by the Ethics Committee of the University of Padua (protocol number: 5326) and performed in accordance with the principles expressed in the 1964 Declaration of Helsinki.

### Procedure

The present study adapted the line bisection task previously used by De Hevia et al, (10) and by De Hevia and Spelke, (11). Before starting the experiment, participants received general instructions about the task. They were informed that on each trial they will be presented with one horizontal line printed on a paper sheet and received instructions to mark with a pencil, rapidly and accurately, the center of each line. They were also told to ignore the flanking stimuli. Stimuli were presented one at a time, aligned with reference to the mid-sagittal plane of the body. The distance between the sheet and the participant was approximately 30 cm. Once the stimulus had been bisected, participants were asked to continue to the next trial.

### Design

The experiments consisted of a baseline phase of 4 trials and a test phase of 128 trials. While in the baseline only the line was displayed, in the test phase the line was presented with flankers on each edge. Flankers in the test phase had two formats, they were either digits (Symbolic format) or dot arrays (Non-Symbolic format). Moreover, when unbalanced, flankers could be oriented in a way that either the most prominent (Large-left) or the least prominent (Small-left) was placed on the left side of the line. This resulted in four conditions (2 formats x 2 orientations) (see Figure 1).

**Figure 1.**
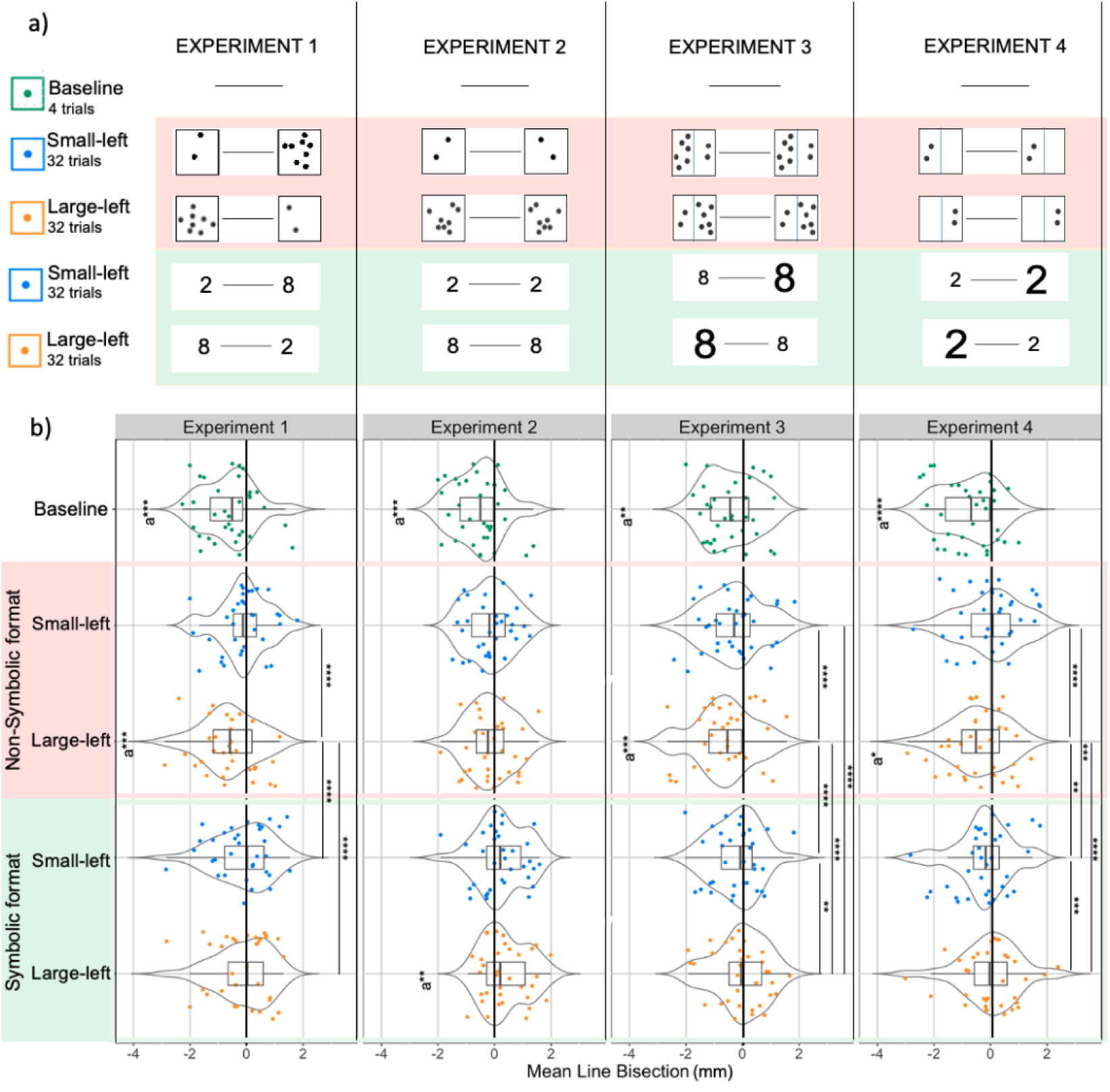
Experimental stimuli and Results of line bisection with Post-hoc analyses in the Linear Mixed-Effects model. In Experiment 1, the flankers were “2-8” or “8-2”. In Experiment 2, the flankers were identical, “2-2” or “8-8”, and symmetrically displaced with respect to the line . In Experiment 3, the stimuli (numerosity 8) were asymmetrically distributed to create a perceptual imbalance. In Experiment 4, we replicated the manipulation used in Experiment 3, but with the numerosity two. All experiments presented the numbers in both symbolic and non-symbolic formats. a) Shows the description of the tasks in the different studies, divided into two blocks: Non-Symbolic (in pink) and Symbolic (in green), with the two orientations Small-Left and Large-Left. b) Shows the results of line bisection with post-hoc analyses in the Linear Mixed-Effects model. The x-axis shows the distance in mm from the centre, for both the Non-Symbolic and Symbolic blocks, each divided into the two orientations Small-Left and Large-Left. The box plots show the mean, standard deviation (ds) and standard error (se), while the violin plots show the distribution of the data with each dot corresponding to one participant. *Note* (* p < .05, ** p < .01, *** p < .001, **** <.0001, a= One Sample t-test against 0)

All the trials of a given condition were presented in 4 blocks with 32 trials each. The order of trials within each block was random. Blocks in the Symbolic and Non-Symbolic format were presented in alternation. The order of presentation was randomized across participants so that half of them completed the Non-Symbolic format first and *viceversa* for the other half.

### Stimuli and Materials

Each participant received a paper block with 132 horizontally oriented sheets (297mm x 105mm). The stimuli were printed in black and white on 12 gsm paper to counteract any interference effects due to the transparency of the sheets. Each sheet depicted a horizontal black line printed in the center. The line was 1 mm wide, the length was either 60 or 80 mm and it was counterbalanced across trials. In the Non-Symbolic format, the stimuli used in Experiments 1 and 2 were generated using the GeNEsIS software (29). The total perimeter was controlled to 7.5 cm for both stimuli. Sixteen different arrays of dots were generated to control and test for configural effects (see <u>OSF</u> https://osf.io/hr8sz/?view_only=bc629be803724e819df7c31e5ddbd213 for major info on each experimental task).

In the Symbolic format, for Experiments 1 and 2, the Arabic digits were printed in a black Arial font, size 30 points (5 mm wide and 7,5 mm high in the printed version). For Experiment 3 and 4, we used two sizes: 30 points and 60 points (10 mm wide and 15 mm high). The digit was horizontally centered and vertically displayed with respect to the invisible square.

In *Experiment 1,* the actual stimuli used were “2” on the left and “8” on the right for the Small-left orientation and *viceversa* for the Large-left orientation (see Figure 1).

In *<u>Experiment 2,</u>* the two flanker stimuli were identical. The stimuli “2-2” were arbitrarily considered the Small-left orientation and the “8-8” were considered the Large-left orientation. The stimuli in the Non-Symbolic format were symmetrical and specular.

In *<u>Experiment 3,</u>* the two flankers were numerically identical (“8-8”), but in an effort to induce a perceptual imbalance while maintaining the same numerosity and configural information, the stimuli were disproportionately distributed. Specifically, in the Non-Symbolic format, a virtual -not visible-vertical line divided the outer frame in two halves with 6 dots in one half and 2 in the other half. In the Small-left orientation, the half with two dots was closer to the left side of the line whereas in the Large-left orientation the half with six dots was closer to the left side of the line.

In *<u>Experiment 4</u>*, like in Experiment 3, the flankers were identical (“2-2”) but unevenly distributed within the stimulus square. More precisely, in the Non-Symbolic format, the flankers were positioned such that the two dots appeared in one half of the square, while the other half was left empty. In the Small-left orientation, the empty half was closer to the left side of the line, whereas in the Large-left orientation the half depicting two dots was displayed to the left side of the line (see Figure 1)

### Data coding

Bisection marks were measured to the nearest mm, rounded to the nearest half unit, using a ruler at the point where the bisection mark intersected the line, as in De Hevia and Spelke (11). Our accuracy using a ruler is approximately ±0.5 mm. Deviations from the objective center towards the left were scored as a negative value, deviations towards the right as a positive value. Participants were excluded from the analyses because their mean bias was more than 2 SD above the group mean.

### Data analysis

In all the studies, Linear Mixed Effects Models (LMMs) were applied to investigate the extent to which participants’ line bisection was influenced by format (Symbolic *vs.* Non-Symbolic) and orientation (Small-left *vs.* Large-left). The dependent variable was the mean deviation in mm of each mark with respect to the objective center. The initial models included the main effects of format and orientation and interactions between these factors. In addition, to control for the repeated measurement in the LMMs, participants were treated as a random factor in all the models. These analyses were performed by fitting a LMMs using the lme4 and lmerTest package (Version 1.1; 30,31) in RStudio (Version 2023.03.1 Build 446; R Development Core Team, 2016). Post hoc tests were conducted by using the emmeans function provided by the emmeans package (32) and the Tukey Method was used for pairwise comparisons while controlling for all family error rates of 4 estimates.

The significance level of statistical tests was predetermined at p = 0.05. Moreover, a one sample t-test was conducted independently of the LMM analysis to evaluate whether the mean deviation in each condition was significantly different from the objective center (zero). This analysis was based on participants’ average responses.

## Results

### Experiment 1

The results showed that flanker format (Symbolic *vs.* Non-Symbolic) had a significant effect (β = 0.46, SE = 0.05, t (4303) = 8.19, p < 0.001), since larger deviations were registered in the Non-Symbolic format (M = −0.35; DS = 1.49) than in the Symbolic format (M = −0.15; DS = 1.62). Orientations (Small-left *vs.* Large-left) had a significant effect on bisection (β = 0.51, SE = 0.055, t(4303) = 9.14, p < 0.001), with larger deviations in the Large-left orientation trials (M =-0.38; DS = 1.59) compared to the Small-left orientation trials (M= −0.12; DS = 1.52).

Additionally, there was a significant interaction between flanker orientation and format (β = −0.50, SE = 0.07, t (4303) = −6.41, p < 0.001) (see Figure 1 and Table in S1).

Post-hoc analyses revealed significant differences between Large-left Non-Symbolic trials and Large-left Symbolic trials (t(4303) = −8.19, p < 0.001) (see all post-hoc tests in S1). Moreover, deviations from the objective center of the line were significant only for the Large-left Non-Symbolic trials (t(33) = −3.63, p < 0.001).

### Experiment 2

The results showed the flanker format (Symbolic *vs.* Non-Symbolic) had a significant impact on bisection (β = 0.50, SE = 0.06, t(4295) = 8.13, p < 0.001), with larger deviations in the Symbolic format (M = 0.30; DS = 1.63) compared with the Non-Symbolic format (M= −0.14; DS = 1.51). Instead, the orientations (Small-left *vs.* Large-left) did not have a significant effect on bisection (β = −0.02, SE = 0.06, t(4295)= −0.40, p = 0.68). The interaction between flanker orientations and format was not significant (β = −0.13, SE= 0.08, t(4295) = −1.46, p = 0.14) (see Figure 1 and Table in S2).

Although the interaction was not significant an exploratory post-hoc analysis was performed. Post-hoc analyses showed significant differences between Large-left Non-Symbolic trials and Large-left Symbolic trials (t(4295) = −8.13, p < 0.001). There were significant differences between Small-left

Non-Symbolic trials and Small-left Symbolic trials (t(4295) = −6.05, p < 0.001) (see all post-hoc tests in S2).

Moreover, deviations from the objective center of the line. The one sample t-test showed significant deviations from zero in Large-left Non Symbolic trials (t(33) = 2.78, p-value < 0.01). There were no other significant differences across conditions.

### Experiment 3

The results showed that the flanker format (Symbolic *vs.* Non-Symbolic) had a significant impact on bisection (β = 0.59, SE = 0.06, t(4297) = 9.72, p < 0.001), with larger deviations in the Non-Symbolic format trials (M = −0.47; DS = 1.68) compared with the Symbolic format trials (M= −0.14; DS = 1.64). Similarly, the orientations (Small-left *vs.* Large-left) had a significant influence (β = 0.31, *SE* = 0.06, t(4297) = 5.17, p < 0.001), with larger deviations in the Large-left orientation trials (M =-0.34; DS = 1.69) compared with the Small-left orientation trials (M= −0.27; DS = 1.63). Notably, an interaction effect was observed between the flanker orientations and format (β = −0.50, *SE* = 0.08, t(4297) = −5.86, p < 0.001) (see Figure 1 and Table in S3).

Post-hoc analyses showed significant differences between Large-left Non-Symbolic trials and Large-left Symbolic trials (t(4297) = −9.72, p < 0.001) (see all post-hoc tests in S3). The one sample t-test results showed significant deviations from zero in Large-left Non-Symbolic trials (t(33) = −3.48, p = 0.001).

### Experiment 4

The results revealed that both flanker format (Symbolic *vs.* Non-Symbolic) and orientations (Small-left *vs.* Large-left) significantly influenced the bisection.

Specifically, there were larger deviations in the Non-Symbolic format trials (M = −0.22; DS = 1.71) compared with the Symbolic format trials (M= −0.12; DS = 1.80) and larger deviations in the Large-left orientations trials (M = −0.23; DS = 1.76) compared with the Small-left orientations trials (M= −0.12; DS = 1.76). Moreover, a significant interaction effect was observed between the flanker orientations and format (β = −0.69, SE = 0.09, t(4304) = −7.62, p < 0.001) (see Figure 1 and Table in S4). Post-hoc analyses revealed significant difference between Large-left Non-Symbolic and Large-left Symbolic (t(4304) = −6.87, p = 0.001) and between Small-left Non-Symbolic and Small-left Symbolic (t(4304)= 3.90, p = 0.0006) (see all post-hoc tests in S4).

The one sample t-test results indicated significant deviations from zero in Large Non-Symbolic trials (t(33) = −2.60, p = 0.01).

## Discussion

In the 4 experiments presented here, a set of numerical stimuli was used to investigate factors that influence the processing of spatial extension. Specifically, our study focused on the line bisection task by exploring how the properties of two flankers positioned at both ends of each line modulated participants’ performance. Crucially, numerical flankers were displayed using both Symbolic (i.e. Arabic numbers) and Non-Symbolic stimuli (i.e. dots). Based on previous studies arguing for the existence of a common system for Symbolic and Non-Symbolic space-number representations (e.g. 11,33,34) we expected that Symbolic stimuli would elicit responses similar to the Non-Symbolic stimuli in this task. Moreover, the physical properties and the numerical value of the flankers were manipulated in an effort to separately explore the relative weight of each of these factors over the orientation of the MNL. Drawing from earlier research, we expected a divergent and opposite bisection in tasks where the small stimulus was on the left compared to tasks where the small stimulus was on the right (5,8,10,11,35).

Not all expectations were confirmed. Most notably, a main effect of the Format was observed across the four experiments, supporting the alternative view that distinct mechanisms might operate in symbolic and non-symbolic processing (20–27). Thus, the present study extends previous literature by providing unprecedented evidence for separate processing in bisection tasks (for a review see 36). Here, differently from previous investigations with the line bisection (e.g. 11) participants systematically tended to show larger leftward biases in the Non-Symbolic format compared to the Symbolic format. The behavior in the Non-symbolic format is compatible with pseudoneglect, namely the phenomenon by which neurologically healthy participants systematically mis-bisect space by erring to the left of the veridical center of a horizontal line (37; consistently observed in the baseline condition across all experiments in our research).

The Symbolic format, on the other hand, seems to entail additional processes that modulate the participants’ behavior. For example, previous studies indicate that Arabic digits increase the saliency of the number magnitude representation in number line tasks via the semantic route (38). Additionally, McCrink and Galamba (34) demonstrated that symbolic numerical information associated with spatial locations improved task accuracy and spatial processing across the visual field. Most relevant for the present findings is the observation that Symbolic processing is strongly left-lateralized in the intraparietal sulcus (39) and it activates the motor areas in the brain more strongly than baseline line bisection tasks without digits (40). This tentatively accounts for the rightward shift in the deviations in the Symbolic format compared to the Non-Symbolic format. We observed those differences, however, only under certain circumstances, as reflected by the interactions between numerical format and orientation. Specifically, the data also showed that the left tendency in the Non-symbolic task was most prominent when the numerically larger flankers were on the left (i.e. Large-left orientation) (Experiment 1) or when the numerically identical flankers with more elements grouped in the proximity of the line were located in the left edge (Experiments 3 and 4), but not when the flankers were numerically equivalent and symmetrical (Experiment 2). By comparing these effects across studies, it seems likely that the imbalance between two flankers (either numerical or perceptual) contributes at directing participants’ attention towards the most salient or larger edge.

It follows that, because our participants in the Non-symbolic format tended to scan first the most salient flanker, their performance in the task varied according to the two orientations presented. In which sense? A previous review and meta-analysis (41) found that participants generally tend to err towards the first cue to which they attended, and therefore, that the direction from which scanning is initiated determines the performance. For example, neurologically normal subjects scanning from left-to-right err significantly to the left of a veridical line midpoint, whereas subjects scanning (with eye or hand) from right-to-left make rightward errors (41). It is thus possible that participants adopt a bisection strategy that systematically scans from left to right, but when oriented in a different direction, results in a different bisection. This is consistent with several studies that manipulated the direction in which lines were visually scanned during a bisection task. The studies found leftward errors in the left-to-right scanning condition and rightward errors in the right-to-left scanning condition (42–45).

This systematically explains our participants’ behavior in the Non-symbolic format. Indeed, the leftward responses consequent of pseudoneglect were enhanced when the prominent flankers were on the left (i.e. Large-left orientation) likely due to a left-to-right scanning over the line. Conversely, when the prominent flankers are on the right (i.e. Small-left orientation), a right-to-left direction bias possibly counteracts the pseudoneglect phenomenon, yielding responses closer to the midpoint. Noticeably, a perceptual imbalance - which even numerically comparable yet asymmetrically distributed arrays can elicit (Experiments 3 & 4) - seems to be at the roots of these effects. Congruently, in the case of perceptual balance (Experiment 2), in which no scanning biases are expected, comparable responses in the two orientations were observed.

The results in the Symbolic format evidence - in agreement with previous research (5,15) - that number and magnitude representations can further modulate the attentional and perceptual effects found for the Non-Symbolic format.

The results in the Symbolic format, specifically in the Large-left orientation, suggest that participants’ attention to the flankers followed the standard numerical order (from smaller to larger). That is, in the case of Large-left orientation trials, they seem to attend the right (Small quantity) first. This prompts rightward deviations that minimize any effects of pseudoneglect, and accounts for the negligible deviations from the veridical center that appear, on average, in these trials. This overall pattern in Large-left orientation trials was found when the symbols differed - and thus could be ordered - on the basis of number (Experiment 1) or physical size (Experiments 3 & 4), but not when they were identical in size and number (Experiment 2). The extensive rightward deviations in the Symbolic format in Experiment 2 reveal how, in the absence of perceptual or numerical imbalance, the mere effects of stimulus format prevail.

It is important to note that this study has some limitations which provide avenues for future research. A first limitation concerns the effect sizes, which appear to be modest and therefore could be influenced by the accuracy of the measurement itself. A further limitation concerns the manipulation of non-symbolic numerosities. It would be beneficial in the future studies to investigate how participants bisect the line when non-numerical variables such as the quantity and area of dots are not congruent with the numerical representations. Finally, to corroborate our interpretations regarding scanning orientations, it would be useful to conduct new research with the use of the eye tracker. This would monitor participants’ eye movements during the experiment, providing a more precise understanding of their visual exploration strategies. Eye tracking could also reveal whether participants explore symbolic and non-symbolic formats differently, providing direct evidence on the role of symbolic numbers in managing attentional resources.

To conclude, the results of these experiments showed distinct effects in line bisection estimations when Symbolic and Non-Symbolic numerosities are used. Specifically, while performance in both Non-Symbolic and Symbolic stimuli is influenced by participants’ perceptual and visuo-spatial attentional biases, only responses in the Symbolic format seem influenced by an organized mental representation of numbers. Therefore, these findings support recent evidence suggesting independent representations for Symbolic and Non-Symbolic numerals (20,24,36,46,47), and with models (48) assuming that the association between numbers and space depends on automatic links but also task-specific features.

## Acknowledgments

**Acknowledgments:** We wish to thank Paola Cazzol and Paria Ahookhosh for their contribution in recruiting participants and data coding of the current paper.

## Supporting Information

**Supporting Information 1.** Linear Mixed-Effects Models Results between flankers Numerosities and Format in Experiment 1.

| Experiment 1 |  |  |  |
| --- | --- | --- | --- |
| Predictors | Estimates | CI | p |
| (Intercept) | -0.60 | -0.90 – -0.31 | <0.001 |
| Orientation [Small-left vs Large-left] | 0.51 | 0.40 – 0.62 | <0.001 |
| Format [Symbolic vs Non-Symbolic] | 0.46 | 0.35 – 0.56 | <0.001 |
| Orientation ×<br>Format | -0.50 | -0.66 – -0.35 | <0.001 |
| Random Effects |  |  |  |
| $\sigma^2$ | 1.68 | | |
| $\tau_{00}$ participant | 0.74 | | |
| ICC | 0.31 |  |  |
| N<sub>participant</sub> | 34 |  |  |
| Observations | 4340 |  |  |
| Marginal R<sup>2</sup> / Conditional R<sup>2</sup> | 0.017 / 0.319 |  |  |

Post-Hoc results in Linear Mixed-Effects Models Results between flankers Numerosities and Format in Experiment 1
| Experiment 1 |  |  |  |  |  |  |  |
| --- | --- | --- | --- | --- | --- | --- | --- |
| *Orientation* | *Format* | *Contrast* | *Emmean* | *Se* | *Df* | *Zratio* | *P.value* |
| Large-left | NON-SYM |  | -0.60 | 0.15 | Inf |  |  |
| Small-left | NON-SYM |  | -0.10 | 0.15 | Inf |  |  |
| Large-left | SYMB |  | -0.15 | 0.15 | Inf |  |  |
| Small-left | SYMB |  | -0.14 | 0.15 | Inf |  |  |
|  |  | (Large-left NON-SYM) - (Small-left NON-SYM) | -0.51 | 0.06 | Inf | -9.14 | **<.0001** |
|  |  | (Large-left NON-SYM) - Large-left SYMB | -0.46 | 0.06 | Inf | -8.20 | **<.0001** |
|  |  | (Large-left NON-SYM) - Small-left SYMB | -0.46 | 0.06 | Inf | -8.28 | **<.0001** |
|  |  | (Small-left NON-SYM) - Large-left SYMB | 0.05 | 0.06 | Inf | 0.95 | 0.78 |
|  |  | (Small-left NON-SYM) - Small-left SYMB | 0.05 | 0.06 | Inf | 0.87 | 0.82 |
|  |  | Large-left SYMB - Small-left SYMB | 0.00 | 0.06 | Inf | -0.08 | 0.99 |

**Supporting Information 2.** Linear Mixed-Effects Models Results between flankers Numerosities and Format in Experiment 2.

**Experiment 2**
| *Predictors* | *Estimates* | *CI* | *p* |
| --- | --- | --- | --- |
| (Intercept) | -0.12 | -0.36 – 0.12 | 0.313 |
| Orientation [Small-left vs Large-left] | -0.02 | -0.14 – 0.09 | 0.683 |
| Format [Symbolic vs Non-Symbolic] | 0.50 | 0.38 – 0.62 | **<0.001** |
| Orientation ×<br>Format | -0.13 | -0.30 – 0.04 | 0.144 |
| **Random Effects** |  |  |  |
| $\sigma^2$ | 2.02 | | |
| $\tau_{00}$ participant | 0.45 | | |
| ICC | 0.18 |  |  |
| $N_{\text{participant}}$ | 34 | | |
| Observations | 4332 |  |  |
| Marginal $R^2$ / Conditional $R^2$ | 0.020 / 0.198 | | |

Post-Hoc results in Linear Mixed-Effects Models Results between flankers Numerosities and Format in Experiment 2
| Experiment 2 |  |  |  |  |  |  |  |
| --- | --- | --- | --- | --- | --- | --- | --- |
| *Orientation* | *Format* | *Contrast* | *Emmean* | *Se* | *Df* | *Zratio* | *P.value* |
| Large-left | NON-SYM |  | -0.12 | 0.12 | Inf |  |  |
| Small-left | NON-SYM |  | -0.14 | 0.12 | Inf |  |  |
| Large-left | SYMB |  | 0.37 | 0.12 | Inf |  |  |
| Small-left | SYMB |  | 0.22 | 0.12 | Inf |  |  |
|  |  | (Large-left NON-SYM) - (Small-left NON-SYM) | 0.02 | 0.06 | Inf | 0.40 | 0.977 |
|  |  | (Large-left NON-SYM) - Large-left SYMB | -0.49 | 0.06 | Inf | -8.13 | <.0001 |
|  |  | (Large-left NON-SYM) - Small-left SYMB | -0.34 | 0.06 | Inf | -5.65 | <.0001 |
|  |  | (Small-left NON-SYM) - Large-left SYMB | -0.52 | 0.06 | Inf | -8.53 | <.0001 |
|  |  | (Small-left NON-SYM) - Small-left SYMB | -0.36 | 0.06 | Inf | -6.05 | **<.0001** |
|  |  | Large-left SYMB - Small-left SYMB | 0.15 | 0.06 | Inf | 2.47 | 0.0638 |

**Supporting Information 3.** Linear Mixed-Effects Models Results between flankers Numerosities and Format in Experiment 3.

| Experiment 3 |  |  |  |
| --- | --- | --- | --- |
| Predictors | Estimates | CI | p |
| (Intercept) | -0.63 | -0.93 – -0.33 | <0.001 |
| Orientation [Small-left vs Large-left] | 0.31 | 0.19 – 0.43 | <0.001 |
| Format [Symbolic vs Non-Symbolic] | 0.59 | 0.47 – 0.71 | <0.001 |
| Orientation ×<br>Format | -0.50 | -0.67 – -0.33 | <0.001 |
| Random Effects |  |  |  |
| $\sigma^2$ | 1.99 | | |
| $\tau_{00}$ participant | 0.74 | | |
| ICC | 0.27 |  |  |
| N participant | 34 |  |  |
| Observations | 4334 |  |  |
| Marginal R<sup>2</sup> / Conditional R<sup>2</sup> | 0.016 / 0.283 |  |  |

Post-Hoc results in Linear Mixed-Effects Models Results between flankers Numerosities and Format in Experiment 3
| Experiment 3 |  |  |  |  |  |  |  |
| --- | --- | --- | --- | --- | --- | --- | --- |
| *Orientation* | *Format* | *Contrast* | *Emmean* | *Se* | *Df* | *Zratio* | *P.value* |
| Large-left | NON-SYM |  | -0.62 | 0.15 | Inf |  |  |
| Small-left | NON-SYM |  | -0.31 | 0.15 | Inf |  |  |
| Large-left | SYMB |  | -0.03 | 0.15 | Inf |  |  |
| Small-left | SYMB |  | -0.22 | 0.15 | Inf |  |  |
|  |  | (Large-left NON-SYM) - (Small-left NON-SYM) | -0.31 | 0.06 | Inf | -5.17 | <.0001 |
|  |  | (Large-left NON-SYM) - Large-left SYMB | -0.58 | 0.06 | Inf | -9.72 | <.0001 |
|  |  | (Large-left NON-SYM) - Small-left SYMB | -0.40 | 0.06 | Inf | -6.60 | <.0001 |
|  |  | (Small-left NON-SYM) - Large-left SYMB | -0.27 | 0.06 | Inf | -4.56 | <.0001 |
|  |  | (Small-left NON-SYM) - Small-left SYMB | -0.08 | 0.06 | Inf | -1.43 | 0.4758 |
|  |  | Large-left SYMB - Small-left SYMB | 0.18 | 0.06 | Inf | 3.12 | **0.0096** |

**Supporting Information 4.** Linear Mixed-Effects Models Results between flankers Numerosities and Format in Experiment 4.

| Experiment 4 |  |  |  |
| --- | --- | --- | --- |
| Predictors | Estimates | CI | p |
| (Intercept) | -0.45 | -0.77 – -0.13 | **0.007** |
| Orientation [Small-left vs Large-left] | 0.45 | 0.33 – 0.58 | **<0.001** |
| Format [Symbolic vs Non-Symbolic] | 0.44 | 0.32 – 0.57 | **<0.001** |
| Orientation ×<br>Format | -0.69 | -0.87 – -0.51 | **<0.001** |
| Random Effects |  |  |  |
| $\sigma^2$ | 2.23 | | |
| $\tau_{00}$ participant | 0.86 | | |
| ICC | 0.28 |  |  |
| N participant | 34 |  |  |
| Observations | 4341 |  |  |
| Marginal R<sup>2</sup> / Conditional R<sup>2</sup> | 0.011 / 0.286 |  |  |

Post-Hoc results in Linear Mixed-Effects Models Results between flankers Numerosities and Format in Experiment 4
| Experiment 4 |  |  |  |  |  |  |  |
| --- | --- | --- | --- | --- | --- | --- | --- |
| *Orientation* | *Format* | *Contrast* | *Emmean* | *Se* | *Df* | *Zratio* | *P.value* |
| Large-left | NON-SYM |  | -0.44 | 0.16 | Inf |  |  |
| Small-left | NON-SYM |  | 0.002 | 0.16 | Inf |  |  |
| Large-left | SYMB |  | -0.007 | 0.16 | Inf |  |  |
| Small-left | SYMB |  | -0.24 | 0.16 | Inf |  |  |
|  |  | (Large-left NON-SYM) - (Small-left NON-SYM) | -0.45 | 0.06 | Inf | -7.03 | <.0001 |
|  |  | (Large-left NON-SYM) - Large-left SYMB | -0.44 | 0.06 | Inf | -6.87 | <.0001 |
|  |  | (Large-left NON-SYM) - Small-left SYMB | -0.20 | 0.06 | Inf | -3.13 | 0.0093 |
|  |  | (Small-left NON-SYM) - Large-left SYMB | 0.009 | 0.06 | Inf | 0.15 | 0.9987 |
|  |  | (Small-left NON-SYM) - Small-left SYMB | 0.24 | 0.06 | Inf | 3.9 | **0.0006** |
|  |  | Large-left SYMB - Small-left SYMB | 0.23 | 0.06 | Inf | 3.74 | **0.001** |

**Supporting Information 5.**
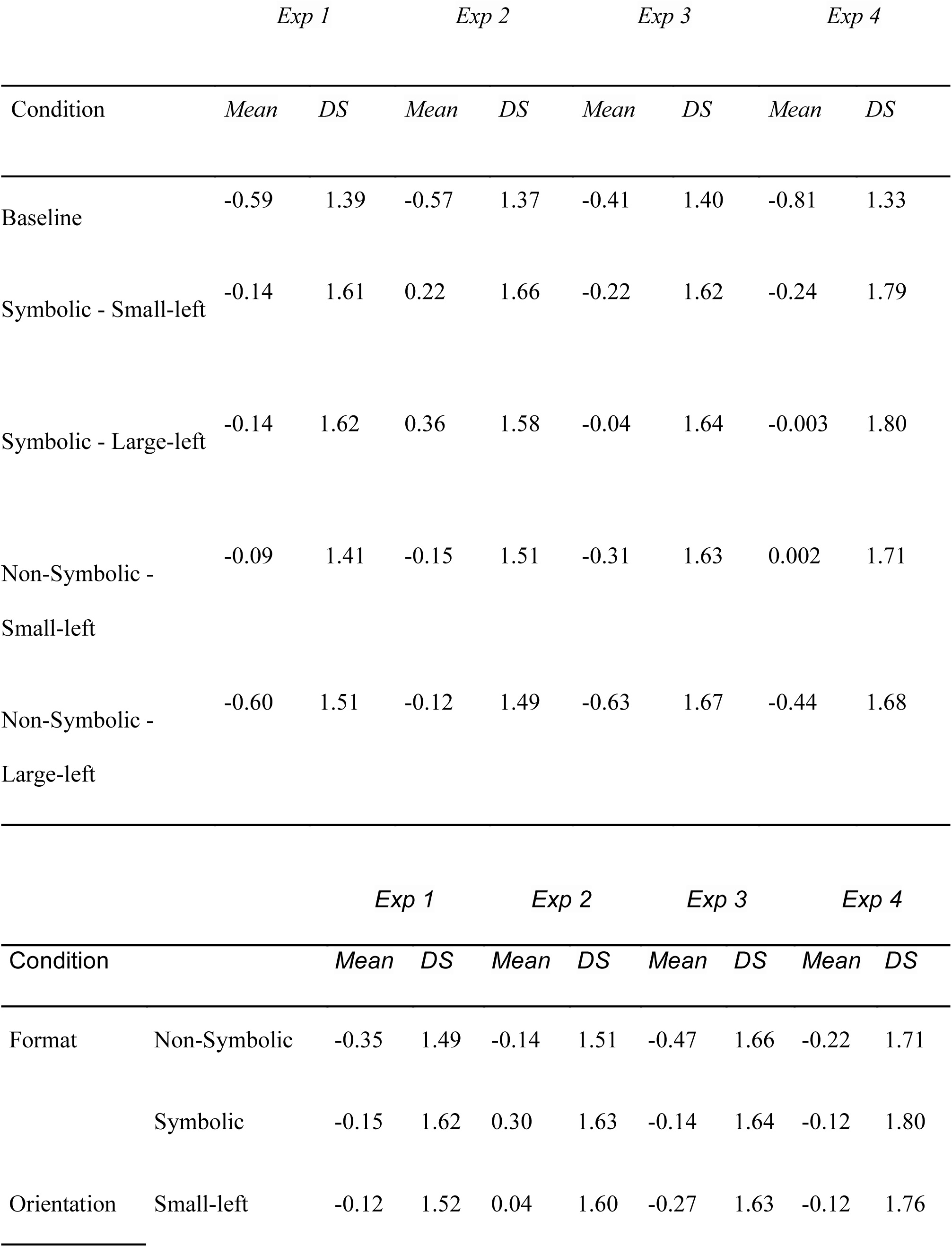
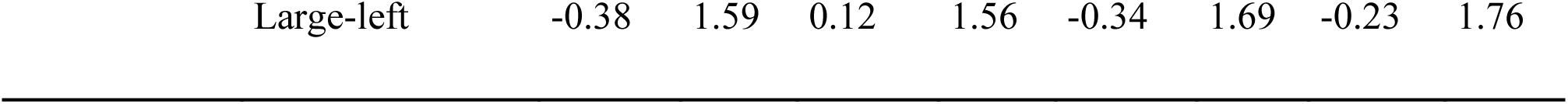
Mean and Standard Deviations of each Experiment.

